# Cell-free Chromatin Immunoprecipitation to detect molecular pathways in Physiological and Disease States

**DOI:** 10.1101/2023.01.24.525414

**Authors:** Moon K. Jang, Tovah E. Markowitz, Temesgen E. Andargie, Zainab Apalara, Skyler Kuhn, Sean Agbor-Enoh

## Abstract

Patient monitoring is a cornerstone in clinical practice to define disease phenotypes and guide clinical management. Unfortunately, this is often reliant on invasive and/or less sensitive methods that do not provide deep phenotype assessments of disease state to guide treatment. This paper examined plasma cell-free DNA chromatin immunoprecipitation sequencing (cfChIP-seq) to define molecular gene sets in physiological and heart transplant patients taking immunosuppression medications. We show cfChIP-seq reliably detect gene signals that correlate with gene expression. In healthy controls and in heart transplant patients, cfChIP-seq reliably detected housekeeping genes. cfChIP-seq identified differential gene signals of the relevant immune and non-immune molecular pathways that were predominantly downregulated in immunosuppressed heart transplant patients compared to healthy controls. cfChIP-seq also identified tissue sources of cfDNA, detecting greater cell-free DNA from cardiac, hematopoietic, and other non-hematopoietic tissues such as the pulmonary, digestive, and neurological tissues in transplant patients than healthy controls. cfChIP-seq gene signals were reproducible between patient populations and blood collection methods. cfChIP-seq may therefore be a reliable approach to provide dynamic assessments of molecular pathways and tissue injury associated to disease.

## Introduction

Defining and monitoring the molecular phenotype of disease guides clinical decision-making including treatment selection, assessment of disease progression and response to treatment. Unfortunately, for many conditions, invasive approaches such as tissue biopsy remain the gold standard. Heart transplant patients, for example, undergo 15 – 25 endomyocardial biopsies to monitor for acute rejection in the first year of transplantation alone. In addition to a high cost and risk of complications, biopsy samples are examined by histopathology, which is limited by low sensitivity and high inter-operator variability (Marboe et al. 2005). Emerging blood-based approaches may address these limitations (Moss et al. 2018; Andargie et al. 2021; Sadeh et al. 2021; Vorperian et al. 2022). Recently, cell-free DNA (cfDNA) chromatin immunoprecipitation sequencing (cfChIP-seq) has been proposed as one of such an approach; being minimally invasive and reliable to define molecular phenotypes and the tissue types involved in multiple disease states (Sadeh et al. 2021). Such an approach could serve as a significant advancement to monitor allograft health. Like other sequencing-based approaches, reproducibility and standardization is important to enable broad applicability and interpretation of cfChIP-Seq results across studies.

Circulating cfDNA, released into the bloodstream following cell death, is attracting great attention as a novel biomarker for early diagnosis and monitoring in a range of disease conditions (Agbor-Enoh et al. 2019; Duvvuri and Lood 2019; Jackson Chornenki et al. 2019; Zviran et al. 2020; Brusca et al. 2022). Given its short half-life in circulation (Lo et al. 1999), and sensitivity, cfDNA analysis mimics tissue/end-organ injury with great temporal precision. Earlier studies to characterize cfDNA focused on genetic differences and DNA methylation-based epigenetic signatures. In transplantation, donor-derived cfDNA (cfDNA), originating from transplanted organs, is a reliable alternative to biopsy. Current approaches use donor-recipient single nucleotide polymorphisms to quantitate donor-derived cfDNA as measure of allograft injury. Although the SNP-based approach is sensitive to detect allograft rejection and other complications, it lacks specificity to identify acute rejection phenotypes or define the molecular pathways involved to tailor treatment. CfDNA is histone bound and maintains histone modifications from its tissue of origin (Sadeh et al. 2021). Post-translational modifications of histones can regulate genomic elements and is often a proxy for gene expression. The associated genes may show cell/tissue specificity, identifying tissue source involved in disease (Sadeh et al. 2021). Thus, cfChip-seq can delineate different disease processes, annotate disease phenotypes, and identify tissue injury patterns (Sadeh et al. 2021).

Despite advances in the development of novel and specific diagnostic approaches, reproducibility remains a major limitation for genomic approaches. In a review of sequence-based studies, only ~25% of the studies provide sufficient information to enable adequate technical and biological reproducibility (Nekrutenko and Taylor 2012). For cfDNA-based studies, blood collection is an added matrix of variability with different collection protocols. Standardization is important to account for and/or limit contamination of cfDNA with cellular genomic DNA from cell lysis during plasma preparation. In one heart transplant study, most of the plasma samples collected were not analyzable because of cfDNA quality, even though specialized blood collection tubes with preservative to prevent blood cell lysis were used (Richmond et al. 2020). Published cfDNA studies continue to use tubes with different preservatives to collect blood and processed blood at different centrifugation speed (Lo et al. 1999; Agbor-Enoh et al. 2017; Sadeh et al. 2021). It remains unknown how these differences contribute to cfDNA results.

Here, in addition to assessing its reproducibility, we assessed if cfChIP-seq can be used to define biological processes in immunosuppressed transplant patients compared to healthy controls, as a first step towards developing specific nucleosome-based cfDNA test that can define molecular states of disease.

## Results

This study intends to assess if a cfChIP-seq approach would delineate biological processes in physiological conditions and, in heart transplantation, as a prototype of disease state. Further, the study assesses the reproducibility of cfChIP-seq between different blood collection methods. Housekeeping genes in healthy controls were selected to reflect physiological conditions. Differential gene signals between immunosuppressed heart transplant patients and healthy controls were performed as a prototype of defined molecular pathways in a disease state. The study involved collection of blood samples and performance of cfChIP-seq on 8 healthy control plasma samples and 6 samples from 2 heart transplant recipients (2 prior to transplantation and 4 post-transplantation). Subjects gave consent. To assess if blood collection methods affect cfChIP-seq results for healthy controls, paired blood samples were collected into two different blood collection tubes; Streck tubes, selected as a prototype sample collection tube containing a proprietary preservative that prevents cell lysis, and EDTA tubes without any anti-cell-lysis preservative. Plasma was separated from blood cells within two hours of blood collection by centrifugation at 1,600 g, followed by centrifugation at two different speeds (3,000 g vs. 16,000 g). So, in total, replicate samples were processed under four conditions: (1) Streck tubes spun at 16,000g (S16), (2) Streck tubes spun at 3000g (S3), (3) EDTA tube spun at 16,000g (E16) and (4) EDTA tubes centrifuged at 3000g (E3) (**Figure 1a**).

**Figure 1:**
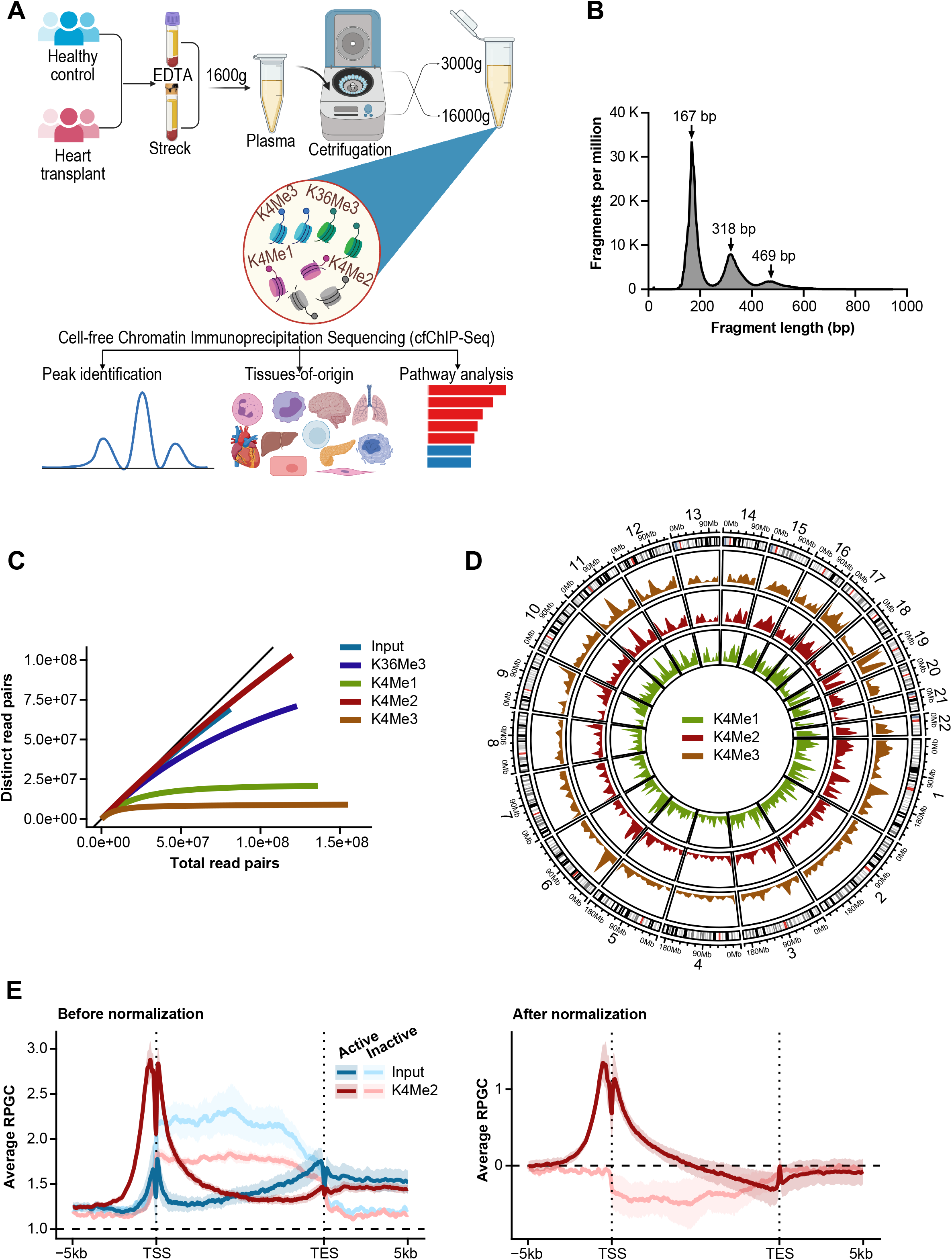
Design of cell-free Chromatin Immunoprecipitation Sequencing (cfChIP-Seq) of plasma samples. **(A)** Schematic workflow: Blood samples of healthy controls and heart transplant patients were collected in EDTA and Streck blood collection tubes. After an initial 1000g centrifugation, plasma was further centrifuged at 3000_g_ or 16000_g_. Cell-free DNA, bound to histones were pooled by specific antibodies which were covalently bound to magnetic beads. Bound cfDNA was purified, used to generate DNA libraries and sequenced. Sequenced reads were analyzed against controls to identify enrichment peaks, tissue-specific signatures and biological pathways. (**B)** Length distribution of sequenced DNA fragments by cfChIP-seq showing nucleosomal periodicity. (**C)** Average sequencing saturation curves for input DNA, K36Me3, K4Me1, K4Me2 and K4Me3 reads, highlighting saturation achieved for H3K4Me3 and H3K4Me2. (**D)** Chromosome-based circos plots representing the distribution of cfChIP-seq peaks across the genome. (**E)** Distribution of cfChIP-seq signals on genes including around the transcription start sites (TSS) and Transcription End Site (TES) for H3K4me2 with and (**F)** without input normalization.

For cfChip-seq, we adopted the protocol from Sadeh et al. (Sadeh et al. 2021). In summary, antibodies directed to specific histone modifications were coupled to magnetic beads and incubated with 1 mL of thawed plasma. After washing and digestion of bound histones using proteinase K, captured cfDNA were indexed using Accel-NGS 2S plus DNA library kit with Unique Dual Indexing (Swift Biosciences). The cfDNA libraries were subject to paired-end sequencing on the NovoSeq platform at ~140 million reads per sample. Two controls were included for each sample, non-specific IgG and input cfDNA. Length distribution of sequence reads showed nucleosomal distribution with mononucleosomal predominance as expected (**Figure 1b**). Of the four histone antibodies, H3K4me2 and H3K4me3 reached saturation at ~30,000 million read pairs per sample; the two other histone antibodies (H3K4me1 and H3K36me3) and input cfDNA did not reach saturation (**Figure 1c, Suppl. Figure 1a**). The three H3K4me marks followed the same global distribution **(Figure 1d)**. Their local distributions matched known patterns, with H3K4me3, for example, found mostly near transcriptional start sites (TSS) (**Figure 1e**), H3K4me2 found at enhancers and TSS sites, and H3K4me1 found primarily at enhancers of housekeeping (active) genes. The number and called peaks for each histone were consistent with these patterns. For example, 46% of H3K4me3 peaks overlapped promoters, compared to 15% for K4Me2 and 18% for K4Me1 **(Suppl. Table 1)**. Use of the input control samples, increased the power of our analyses. It increased the number of peaks identified for all histone modifications **(Supp. Table 1)**. It also corrected for a weak binding pattern seen on gene bodies for non-expressed genes for all histones **(Figure 1e, Suppl. Fig 1b)**. For these reasons, input-normalization was utilized for all downstream analyses.

### Reproducibility of cfChIP-seq signals across blood collection methods

We first evaluated if the four different blood collection and processing conditions alter cfChIP gene signals for H3K4me3, H3K4me1, H3K4me2, and H3K36me3. All four conditions, i.e S3, S16, E3, and E16, showed expected peak frequency distribution for H3K4me3 around the TSS for expressed genes consistent with known promoter location (**Figure 2a**). Expected frequency distribution of gene signals was also observed for H3K4me1, H3K4me2, and H3K36me3 (**Suppl. Figure 2**). For all four blood processing conditions, FRiP were highest for H3K4me3 than for H3K4me1 and H3K4me2 (**Figure 2b**). Fraction of reads in peaks (FRiP) was lower and/or more variable between subjects for E16 and E3 than for S16 and S3; E3 showing the most variable FRiP, while S3 and S16 showed more consistent FRiP between subjects for H3K4me1, H3K4me2, and H3K4me3 (**Figure 2b**). The decrease in FRiP in EDTA tubes compared to Streck tubes was significant (p-value: 2e^−4^). The number of peaks detected was equally variable for E3, E16 than for S3 and S16 (**Figure 2c**). Despite the variability in peaks between the conditions, for all four conditions, H3K4me3 cfChIP-seq showed similar fraction of promoters overlapping peaks (**Figure 2d**), as well as a similar number of genes detected (**Suppl. Table 1)**. H3K4me1 and H3K4me2 also showed similar number of genes detected, except for one sample that showed lower number of genes detected for E3 (**Suppl. Table 1**).

**Figure 2:**
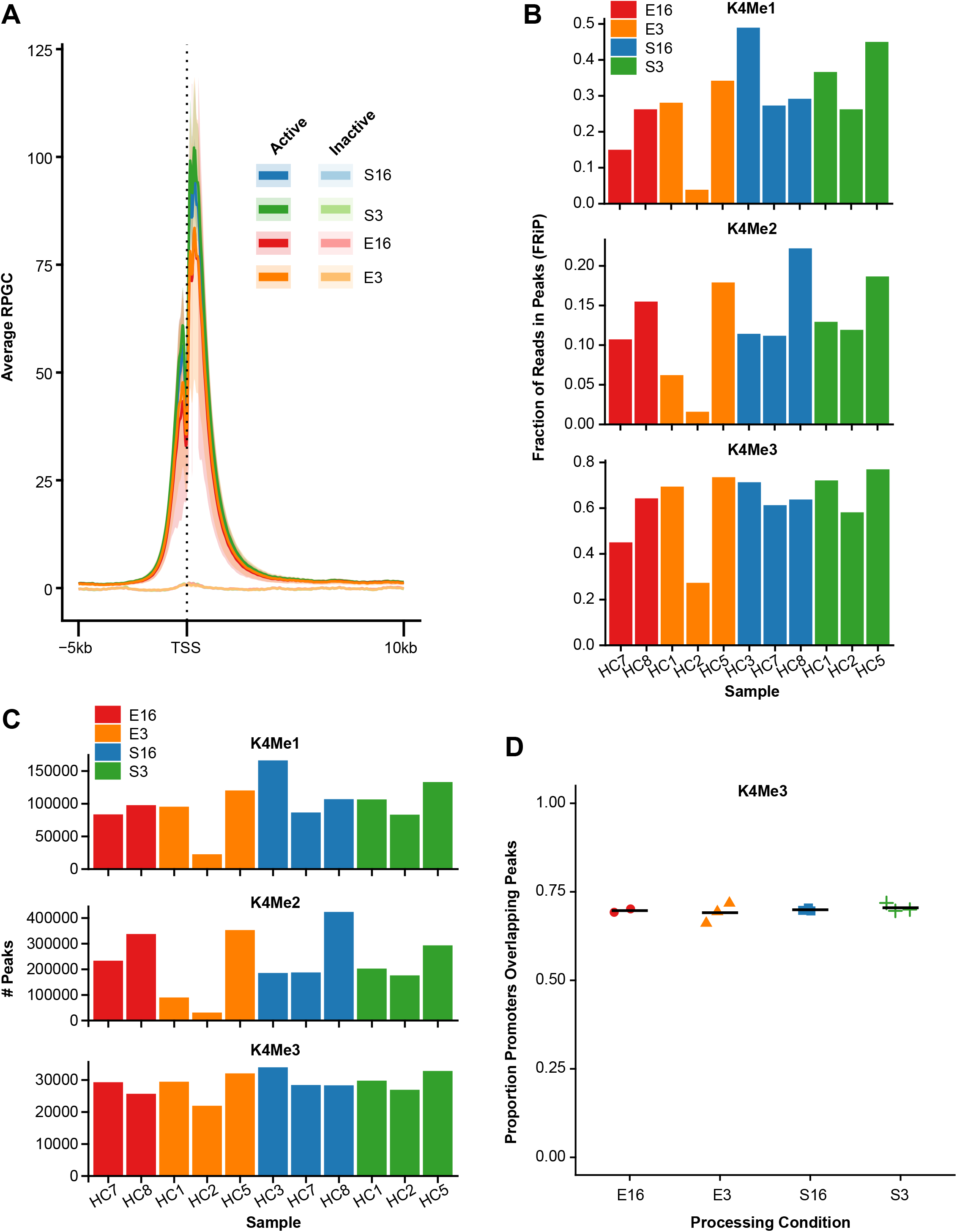
Assessing replication of cfChIP-seq signals across different sample collection and processing conditions. **(A)** K4Me3 distribution around the TSS in different processing conditions: E16 [red], E3 [orange], S16 [blue], S3 [green]. **(B)** Bar plots showing Fraction of Reads in Peaks (FRiP) for K4Me1 (left), K4Me2 (middle) and K4Me3 (right) across the four blood processing conditions. Each bar on the x-axis indicates individual subjects and the color indicates the processing conditions. FRiP were highest for H3K4me3 than for H3K4me1 and H3K4me2. **(C)** Bar plots representing the number of called peaks of individual samples for K4Me1 (top), K4Me2 (middle) and K4Me3 (bottom) across the four blood processing conditions. **(D)** The fraction of protein-coding gene promoters overlapping K4Me3 peaks across the four blood processing conditions.

### CfChIP-seq reliably detect housekeeping genes

We next assessed if cfChIP-seq gene signals are reflective of gene expression, focusing on S16/H3K4me3, which showed highest and most consistent RPGC signal between subjects. Prototype housekeeping genes (GAPDH and TBP) show peaks matching their promoter location, around the TSS. Similarly, example monocyte-specific genes (FCN1 and CSF3R), a third major cell type contributing plasma cfDNA, showed peaks matching their gene promoter location. However, the monocyte-specific genes showed lower RPGC compared to housekeeping genes that are constitutionally expressed in all tissues. Non-expressed genes in healthy patients (IL-3 and CSF2) showed no or non-specific peaks, with baseline levels that are no different from non-specific IgG (**Figure 3a**). In total, H3K4me3 cfChIP-seq detected 93% of housekeeping genes, (**Figure 3b**), that is 8486 of the 9099 housekeeping genes represented in **Suppl. Table 2**. Of the 4216 non-housekeeping genes detected by H3K4me3 cfChIP-seq, one-third (n=1475 genes) were neutrophil and/or monocyte-specific genes; neutrophils and monocytes contribute over one-third of plasma cfDNA (Moss et al. 2018; Andargie et al. 2021) (**Figure 3b**). Since leukocytes contribute over three-quarters of plasma cfDNA in healthy patients (Moss et al. 2018), we determined if K4Me3 cfChIP-seq gene signals correlate with leukocyte ChIP-seq, and observed a strong correlation (**Figure 3c**). Taken together, these findings are consistent with the prior report (Sadeh et al. 2021) and indicate that H3K4me3 cfChIP-seq reliably detect gene expression signals that are biologically plausible in healthy controls.

**Figure 3:**
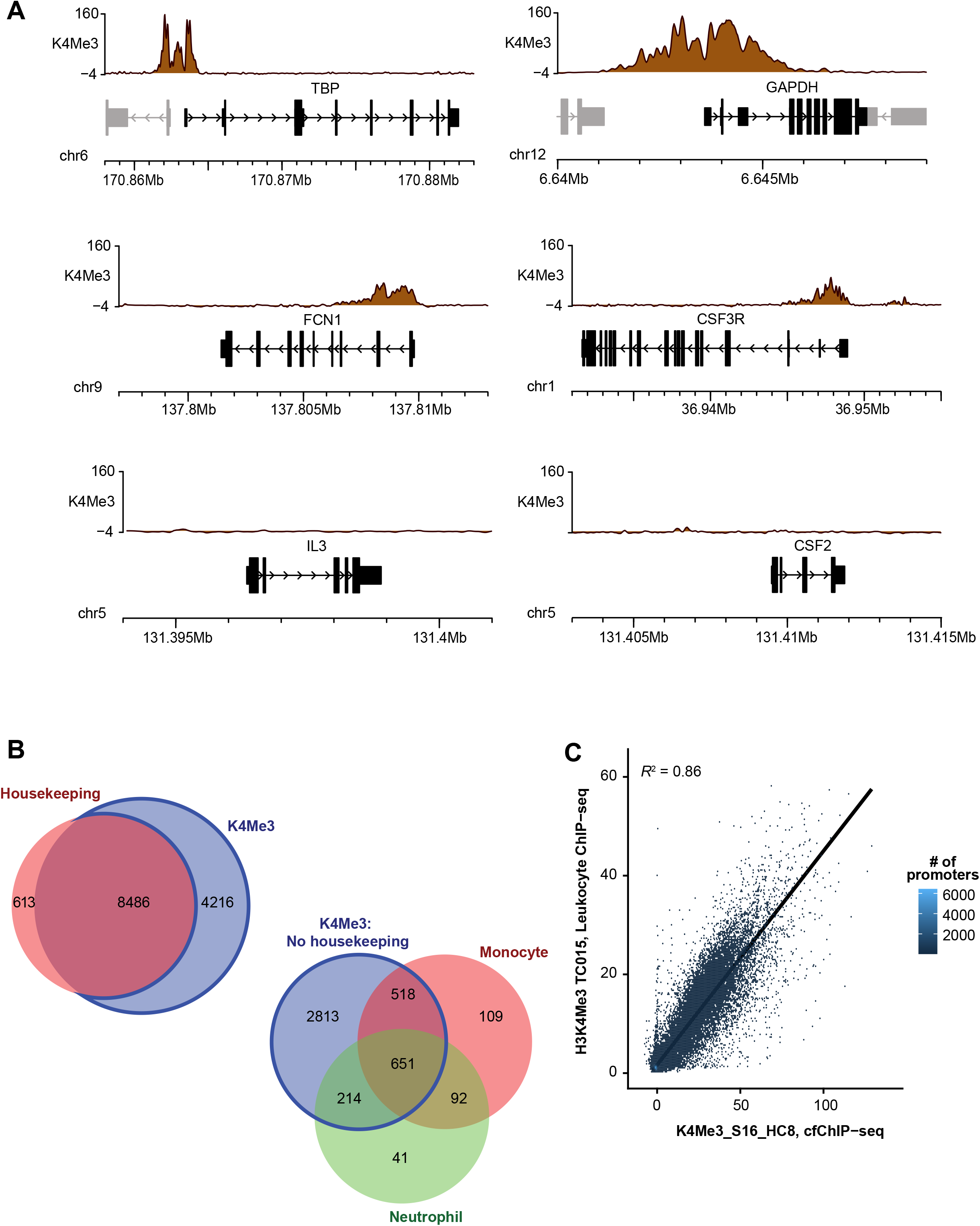
ChIP-Seq correlates with gene expression in physiological state. **(A)** Epigenome browser snapshots showing (RPGC-normalized sequence reads) H3K4me3 ChIP-seq signals at example housekeeping (TBP, GAPDH), monocyte-specific genes (FCN1, CSF3R) and silent (IL-3, CSF2) genes. **(B)** Comparison of genes identified by ChIP-Seq. Top: Venn diagram showing the number of promoters overlapping H3K4Me3 peaks that are housekeeping genes(red). All gene lists were defined by (Sadeh et al. 2021) and identified in Supplemental Table 2. Bottom: Venn diagram showing the overlap of non-housekeeping genes having H3K4Me3 peaks associated with monocytes (orange) and/or neutrophils (green). **(C)** Scatterplot plot showing the correlation between H3K4Me3 leukocyte ChIP-seq data and cfChIP-seq data (R^2^ = 0.86). Leukocyte data from (Kundaje et al. 2015).

### CfChIP-seq identifies relevant molecular pathways in heart transplant patients

We next analyzed plasma from heart transplant recipients maintained on immunosuppression drugs. The patients involved all reached theurapeutic tacrolimus blood toughs by 2 weeks of transplantation. So, post-transplant blood samples were collected 28 – 318 days after transplantation. Blood samples were collected in Streck tubes and spun to 1,600g followed by 16,000g. The isolated cfDNA showed an expected nucleosomal pattern (**Suppl. Figure 3a**). H3K4me3 peaks showed highest frequency around TSS as expected (**Suppl. Figure 3b**). Like for healthy controls, cfChIP-seq reliably detected housekeeping genes in heart transplant patients, detecting 8511 of the 9099 housekeeping genes. The remaining 3872 non-housekeeping genes detected were predominantly associated to monocytes and neutrophils (n=1483) (**Figure 4a)**. A principal component analysis (PCA) showed separation of cfChIP peaks of healthy controls and transplant recipients. To verify that the PCA separation was due to difference in gene expression signals and not due to different in the healthy control and heart transplant groups, we included two pre-transplant plasma samples collected from the two heart transplant recipients. Both heart transplant recipients had advanced dilated cardiomyopathy and were status 1A on the transplant wait list at the time of sample collection. PCA showed separation of cfChIP-seq signals for the pre-transplant samples versus healthy controls, as well as separation of signals for the pre-transplant and post-transplant samples **(Figure 4b**).

**Figure 4:**
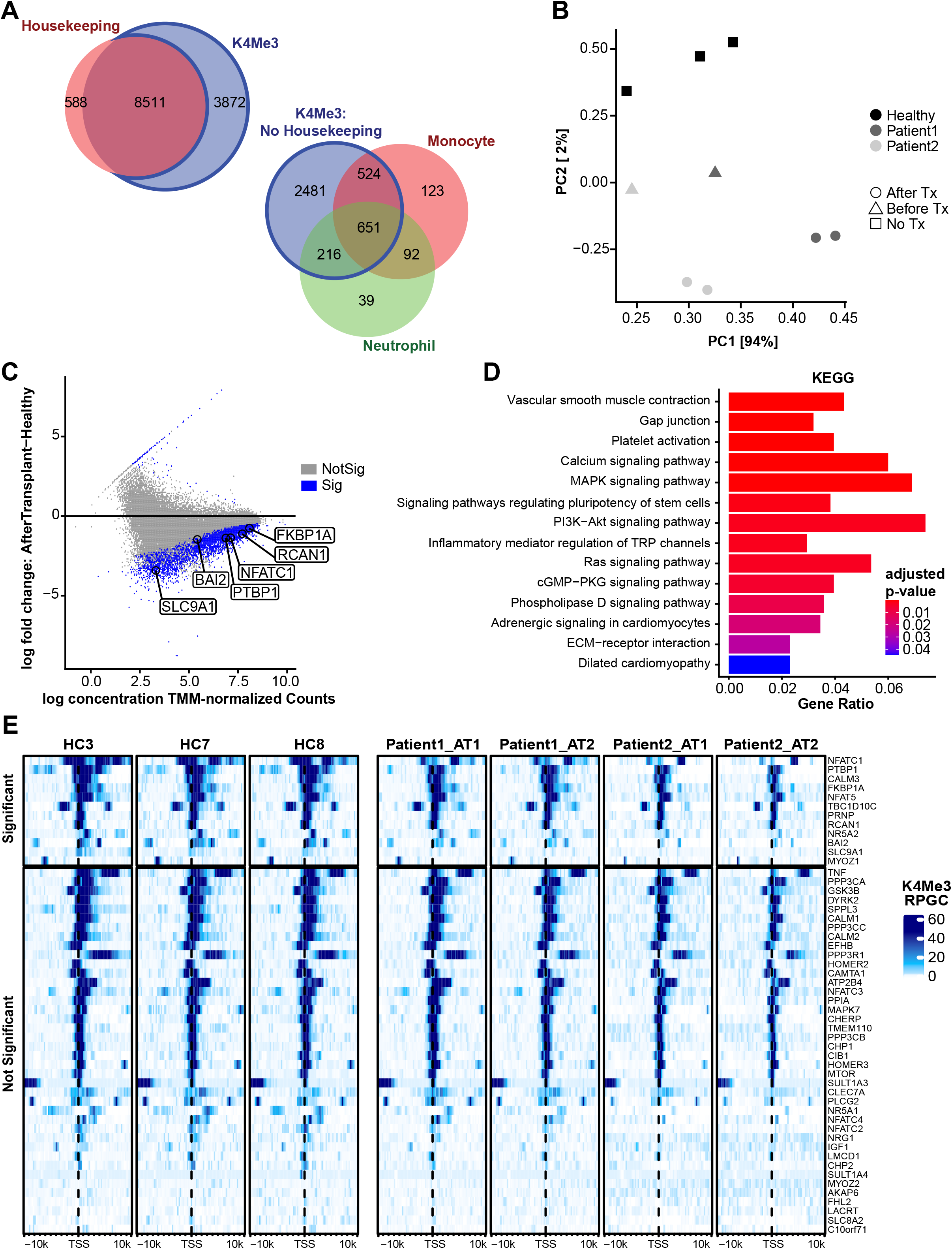
H3K4Me3 cfChIP-seq identifies relevant genes in heart transplant patients. **(A)** Venn diagram showing H3K4Me3 peaks in transplant patients overlapping promoters of constitutively active housekeeping genes, as well as non-housekeeping genes associated to monocytes (top) and and neutrophils (bottom). **(B)** Principal component analysis (PCA) plot of H3K4Me3 peaks showing separation of healthy controls from pre-transplant and post-transplant patient samples **(C)** MA scatter plot shows differential gene peaks between heart transplant and healthy controls; a subset of calcineurin genes are marked. The blue dots indicate significant differential gene signals between heart transplant and healthy controls, marked dots under the thick line depict genes with lower H3K4Me3 signals in heart transplant subjects compared to healthy controls. **(D)** KEGG pathway enrichment analysis of genes whose promoters were associated with a significant negative fold-change in transplant patients relative to healthy controls. Select immune and non-immune pathways associated with transplantation state are shown. **(E)** Heatmap showing H3K4Me3 pattern around the TSS site of genes associated with the calcineurin pathway. Calcineurin genes were defined as being members of GO:0097720 or Reactome R-HSA-2025928. Genes above the thick line shows significant difference specific to transplantation state as compared to healthy controls.

We employed the Deseq2 version of DiffBind v2 (Ross-Innes et al. 2012) to identify differential peaks between healthy controls and heart transplant recipients. There were 4030 peaks with a FDR less than 0.05 between the healthy controls and heart transplant samples (**Figure 4c)**. 48.8% of these peaks overlapped promoters of 2137 genes. Pathway analysis of these genes identified immune and non-immune pathways that are relevant to transplantation (**Figure 4d, Suppl. Table 3**). The immune pathways were predominantly associated with a loss of H3K4Me3 in their promoters in transplant recipients versus healthy controls, correlating with the immunosuppressed state of these transplant patients. Indeed, blood tacrolimus trough at time of sample collection was 8 – 12 ng/dL in these patients and expected to be zero for healthy controls who are not on tacrolimus. Of the immune pathways identified, multiple genes showed the differential pattern, including 55 genes in the Rap1 pathway and 42 genes in the Ras signaling pathways. Given that heart transplant patients are maintained on tacrolimus, a calcineurin inhibitor, we further analyzed calcineurin pathway genes. NFACT1, the main target of calcineurin inhibitors, as was 11 of the other 51 calcineurin related genes showed differential cfChIP-seq signals between transplant patients and healthy controls (**Figure 4f**). Non-immune pathways also showed differential cfChIP-seq signals between transplant patients and healthy controls (**Suppl. Table 3)**. Genes associated to re-organization of extracellular matrix and fibrosis showed differences between transplant and healthy controls, including genes associated to glycosaminoglycan metabolism (n=26), extracellular matrix organization (n=57), collagen synthesis (n=21), and integrin synthesis/re-organization (**Suppl. Table 3)**. Interesting, multiple gene sets associated to neuropathy were also differentially detected in transplant patients, neuropathy due to drug toxicity is a common manifestation in heart transplant patients.

### Different cfDNA tissue sources in transplant patients and healthy controls

CfDNA was assigned to different tissue types using a library of tissue-specific genes signals of 27 tissues (Sadeh et al. 2021). In healthy controls and transplant patients, hematopoietic cells were major contributors of cfDNA (**Figure 5a**), contributing 85% of cfDNA (**Figure 5b**). Comparing the four blood processing methods, S16 and S3 produced more consistent cfDNA tissue contributions than E16 or E3 (**Suppl. Fig 4a**). Vascular endothelial cells, pulmonary tissues and other non-hematopoietic tissue types were also detected. The fraction of tissue-specific cfDNA detected by cfChIP-seq correlated with tissue contributions determined using bisulfite sequencing (**Figure 5C**) using data from a prior publication. The latter approach uses tissue-specific DNA methylation signatures to assign tissue-specific cfDNA. Quantitatively, total cfDNA was 10 times higher in transplant patients than healthy controls (**Figure 5d**). Cardiac-specific cfDNA was 6 times higher in the heart transplant patients than healthy controls (**Figure 5e**), expected in these patients where the heart allograft is exposed to host immunity directed against the allograft. To further assess the greater heart injury in transplant patients, we measured allograft or donor-derived cfDNA (ddcfDNA) by digital droplet PCR using primers directed to donor-recipient single nucleotide polymorphisms. Again, we observed higher levels of ddcfDNA fraction (0.45%) in transplant patients, compared to healthy controls who showed levels similar to background (<0.01%). In addition, transplant patients showed higher tissue-specific cfDNA from hematopoietic and other non-hematopoietic tissues (**Figure 5 F & G**). For example, gastro-intestinal-specific cfDNA, and neuron-specific cfDNA were 15 and 20 times higher in transplant patients compared to healthy controls (**Figure 5G)**, which correlate with the increased frequency of gastrointestinal or neuropathy symptomatology (Díaz et al. 2007; Şahintürk et al. 2021) observed in transplant patients. Taken together, cfChIP-seq detects biologically plausible gene sets and pathways in heathy controls and heart transplant patients, demonstrating adequate reproducibility with different blood collection protocols. In addition, cfDNA identifies the relevant tissue involvement in heart transplant patients.

**Figure 5:**
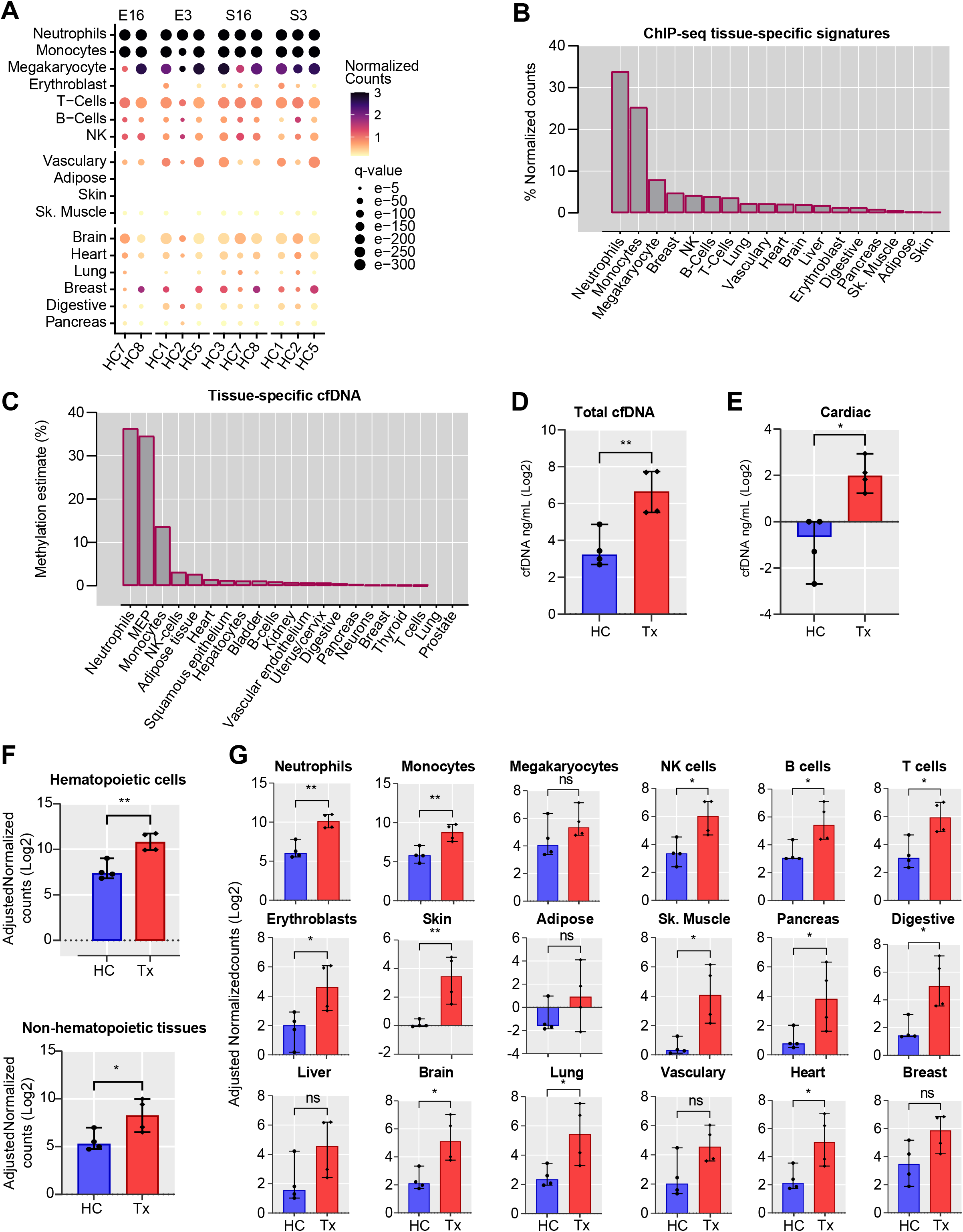
cfChIP-seq identifies tissue sources of cfDNA. (A) Dot plot showing the cell or tissue contribution of plasma cfDNA in healthy controls estimated. The radius of the circle represents Benjamin–Hochberg-adjusted p-values (q-scores) and the color represents the magnitude of normalized reads per kilobase (kb). **(B)** Bar plot showing the proportion of cfChIP-seq signatures from different tissue types, calculated from the normalized counts **(C)** Bar plot showing the distribution of plasma cell or tissue-specific cfDNA level using whole-genome bisulfite sequencing (WGBS) (Andargie et al. 2021). **(D)** The concentration of total plasma cfDNA (ng/mL) was determined by qPCR in healthy controls and transplant patients. **(E)** Comparison of cardiac-specific cfDNA in healthy controls and heart transplant patients, measured using cfChIP-seq. Absolute concentration was measured by multiplying cardiac-specific estimate (%) by total cfDNA concentration. **(F)** Comparison of cfChIP-seq hematopoietic- and non-hematopoietic tissue signatures in healthy controls and heart transplant patients. **(G)** Comparison of cfChIP-seq tissue-specific signatures in healthy controls and heart transplant patients. For plots G-U, input normalized reads/kb was multiplied with total cfDNA concentration and comparison was done by Mann Whitney U test. P-value <0.05 was considered statistically significant; *p < 0.05; **p < 0.01; ***p < 0.001.

## Discussion

In disease management, molecular of clinical and often molecular indicators is often needed to guide treatment selection or monitor treatment response. Unfortunately, this remains a limitation for many disease conditions such as transplantation where the existing tools are invasive and/or have poor sensitivity (Marboe et al. 2005). This study shows that plasma cfDNA chromatin immunoprecipitation sequencing (cfChIP-seq) may provide a minimally invasive approach to identify biologically plausible gene and molecular pathway signals in both heathy controls and heart transplant patients. The approach also profiles tissue-specific cfDNA and thus, identifies tissue injury patterns that are biological plausible. We further demonstrate that cfChIP-seq approach provides replicable gene signals with different blood collection methods The study findings are potentially broad applicability concurrent with a prior study.

cfChIP-seq is a proxy of gene expression, with over 90% of housekeeping genes consistently identified in healthy control subjects and heart transplant recipients. The non-housekeeping genes identified were predominantly from cells that are major contributors of plasma cfDNA (Sadeh et al. 2021). In the prototype cardiac transplant recipients, many immune-related gene signals were downregulated compared to healthy controls. This is expected in these patients who are maintained on multiple therapies for immunosuppression. The immune pathways identified are relevant in transplantation (Fric et al. 2014; Wang et al. 2015; Bonezi et al. 2020). Indeed, several genes of the calcineurin pathways were down regulated in these patients who are maintained on calcineurin inhibitors. In addition to identifying relevant immune pathways, cfChIP-seq identified differential gene signals related to extracellular matrix re-organization and fibrosis (Chandra et al. 2021; Fan and Hu 2022). Allograft fibrosis is typically observed 5 – 15 years post-transplantation when long-term complications and chronic rejection involving these pathways develop. Additional studies are needed to define the relationship of these early gene changes to chronic allograft fibrosis and chronic rejection, two long-term sequalae that involve extracellular matrix remodeling.

The cfChIP-seq approach also identified tissue contributions of cfDNA that were consistent in prior report (Sadeh et al. 2021). Heart transplant patients show higher cardiac specific-cfDNA compared to healthy controls, expected in these patients. The higher cardiac injury correlated with high allograft-derived cfDNA. We also observed elevated cfDNA from multiple hematopoietic and non-hematopoietic tissue types. The tissue-injury pattern observed in heart transplant patients is biological plausible and represent tissue types with increased incidence of symptomatology observed in these patients due to drug toxicity(Lindenfeld et al. 2004; Tayyem et al. 2018), infections or other complications (Díaz et al. 2007; De Weerdt et al. 2008; Pruitt et al. 2013; Şahintürk et al. 2021).

Studies with larger number of subjects will be relevant to validate this study findings and further assess the clinical utility. These future studies should sequence cfDNA to higher sequencing depth to fully assess the utility of cfChIP-seq signals for H3K4me1, H3K36me3 and other histone antibodies. Studies assessing cfChIP-seq reproducibility following different sample storage conditions are also needed. Pending these additional studies, this pilot study indicates that cfChIP-seq reliable defines gene expression signals that are relevant in heart transplantation. Such a minimally invasive approach may be utilized to reliable monitor transplant patients and other patient population.

## Methods

### Patient recruitment

Healthy adult control plasma samples were obtained at the time of blood donation at the NIH Clinic Center as part of a Department of Transfusion Medicine protocol (99-CC-0168; Collection and Distribution of Blood Component from Healthy Donors for In Vitro Research Use; ClinicalTrials.gov NCT00001846). The protocol uses a questionnaire-based approach to rule out any known chronic disease in healthy controls. Heart transplant patients were recruited as part of the Genome Transplant Dynamics, a multicenter study supported by the Genomic Research Alliance for Transplantation (GRAfT, ClinicalTrials.gov NCT02423070). The study consent and recruit patients awaiting heart transplantation and monitor prospectively after transplant with collection of plasma samples and clinical data. All subjects provided written informed consent.

### Blood collection and processing protocols

For each healthy control patient, two 8 – 10 mL blood samples were collected simultaneously into Streck cell-free DNA BCT tube (Streck) or into BD Vacutainer plastic EDTA tube to prepare plasma. All peripheral blood samples were first centrifuged at 1,600 g for 10 min at 4 °C. Plasma samples for half of the patients (both Streck and EDTA) underwent a second centrifugation at 3,000 g or 16,000 g for 10 min at 4 °C. One milliliter aliquots of plasma was stored at −80 °C until use. For transplant patients, blood was collected into Streck tubes and centrifuged at 1,600 g followed by a 16,000 g. One milliliter aliquots of plasma was stored at −80 °C until use.

### cfDNA chromatin immunoprecipitation and library construction

The cfDNA chromatin immunoprecipitation and library construction were performed as described (Sadeh et al. 2021), with minor modifications. In brief, we performed ChIP reactions with covalently conjugated antibodies to magnetic beads to improve the efficiency of ChIPed cfDNA. Fifty micrograms of antibody were conjugated to 5 mg of epoxy M270 Dynabeads (Invitrogen) according to manufacturer instructions. The antibody conjugated beads were stored at 4 °C in PBS with 0.02% sodium azide solution. After washing once, 20 uL (0.2 mg, ~2 μg of antibody) of conjugated beads plus 0.1% BSA, 1 mL of plasma was directly added to the rinsed beads in a deep well plate on magnetic fields. In addition, 1X protease inhibitor cocktail (Roche) and 10 mM EDTA was added, and the reaction was mixed by rotating overnight at 4 °C. The beads were magnetized and washed 8 times with 150 μl of blood wash buffer (50 mM Tris-HCl, 150 mM NaCl, 1% Triton X-100, 0.1% sodium deoxycholate, 2 mM EDTA, 1× protease inhibitor cocktail) on ice and followed by washing 3 times with 150 μl of 10 mM Tris pH 7.4 on ice. The histone bound beads were resuspended and the histones were eluted by incubating for 1 h at 55 °C in 50 μl of chromatin elution buffer (10 mM Tris pH 8.0, 5 mM EDTA, 300 mM NaCl, 0.6% SDS) containing 50 units of proteinase K (Epicenter). The ChIPed cfDNA were purified by 1.4X SPRI cleanup (AMPure XP, Agencourt) and processed for DNA library construction by indexing on beads using Accel-NGS 2S plus DNA library kit (IDT).

### Extraction of Input cfDNA and NGS sequencing

The volumes of plasma were adjusted to 2.1 mL with PBS containing certain concentration of lambda (λ) DNA shared to ~170 bp to make the final concentration to 0.143 ng/mL plasma. cfDNA was isolated by QIAsymphony circulating DNA kit using a customized 2 mL protocol in 60 uL elution volume. The isolated cfDNA was quantified by QPCR with human Alu115 primers (Forward (F’)- CCTGAGGTCAGGAGTTCGAG/Reverse (R’)-CCCGAGTAGCTGGGATTACA) and Alu247 primers (F’-GTGGCTCACGCCTGTAATC/R’-CAGGCTGGAGTGCAGTGG). The integrity of cfDNA was estimated as the Alu247/Alu115 ratio and the recovery rate calculated with the eluted/added ratio of λ-DNA by QPCR with specific λDNA primers (F’- CGGCGTCAAAAAGAACTTCC/R’- GCATCCTGAATGCAGCCATA). To prepare DNA library for input cfDNA, Accel-NGS 2S plus DNA library kit was used with 1 ng of cfDNA with 9 cylces of PCR amplification. The indexed DNA library was sequenced on Novaseq6000 as following standard procedures (~1.4×10^8^ properly aligned reads, 2X100 bp).

### ChIP-seq processing

Reads were trimmed with Cutadapt version 1.18 (Kechin et al. 2017). All reads aligning to the Encode hg19 v1 blacklist regions (Moore et al. 2020) were identified by alignment with BWA version 0.7.17 (Li and Durbin 2009) and removed with Picard SamToFastq (**https://broadinstitute.github.io/picard/)**. Remaining reads were aligned to an hg19 reference genome using BWA. Reads with a mapQ score less than 6 were removed with SAMtools version 1.6 (Li et al. 2009) and PCR duplicates were removed with Picard MarkDuplicates. Data was converted into bigwigs for viewing and normalized by reads per genomic content (RPGC) using deepTools version 3.0.1 (Ramírez et al. 2016) using the following parameters: --binSize 25 -- smoothLength 75 --effectiveGenomeSize 2700000000 --centerReads --normalizeUsing RPGC. RPGC-normalized input values were subtracted from RPGC-normalized ChIP values of matching cell type genome-wide using deepTools with --binSize 25.

### Peak calling and annotation

Prior to peak calling, reads aligning to chromosomes X, Y, and M were filtered using SAMtools. Peaks were called using macsNarrow (macs version 2.1.1 from 2016/03/09) (Zhang et al. 2008) with the following parameters: −q 0.01 --keep-dup=“all” −f “BAMPE”. Differential peaks were called using DiffBind v2 and its Deseq2 differential caller with default parameters. Peaks were annotated using UROPA version 4.0.2 (Kondili et al. 2017) and Gencode Release 19 (GrCh37).

UROPA annotation conditions involved three query steps, each having an attribute.value filter of “protein_coding” and a feature.anchor of “start”. The three queries varied only by distance which were set as: 3000, 10000, and 100000. For gene annotations, only the closest match (finalhit) was used for downstream analyses. For promoter annotations, all genes that matched the first query (allhit) were used for downstream analyses. Over-enrichment analysis was completed on the promoter annotations for H3K4Me3 using clusterProfiler and plotted with enrichplot. Specifically, the list of genes was compared to the KEGG human pathways with an appropriate background set.

### Statistical analysis and visualization

Graphs were generated by the R software (v3.6.3 or later) using Ggprism, Ggplot, Ggrepel, patchwork, VennDiagram, EnrichedHeatmap, circlize, and karyoploteR. Python software (v3.5) using pybedtools and pysam packages and GraphPad Prism software (v9.4.1) were also used. Comparisons between groups were performed using nonparametric Mann-Whitney U test either on GraphPad Prism or R software (v4.0.3) and adjusted for multiple testing using the Bonferroni correction. P < 0.05 indicates statistically significant: *P < 0.05, **P < 0.01, ***P < 0.001, and ****P < 0.0001. For the FRiP statistical test, differences FRiP between conditions were modeled using linear mixed effects regression with no fixed effects (intercept-only) and a random effect for bam files within each sample. A t-test was performed to assess statistical significance using the nlme package in R.

## Competing interest

All the authors have no conflicts of interest to declare.

## Acknowledgments

This work was supported, in part, by intramural research funds of the National Heart, Lung, and Blood Institute (NHLBI) and Lasker Clinical Research Fellowship Program. The authors thank Dr. Randall Johnson for assistance in statistical analyses.

## Author contributions

M.J.K. designed and performed experiments, analyzed data, and wrote the paper. T.E.M performed bioinformatics analysis and wrote the paper. T.E.A. performed experiments and analyzed data and wrote the paper. Z.A. performed bioinformatics analysis. S.K. performed bioinformatics analysis. S.E. conceived the original idea, supervised the study, and wrote the paper with input from all authors.

## Data and code availability

The authors declare that all data supporting the findings of this work are available within the article and its Suppl. information, and raw sequencing data are available from the corresponding author upon reasonable request. All codes for cfChIP-seq data analysis is available at github repository: https://github.com/OpenOmics/cfChIP-seek.

## Supplementary Figure Legend

**Suppl Figure 1.**
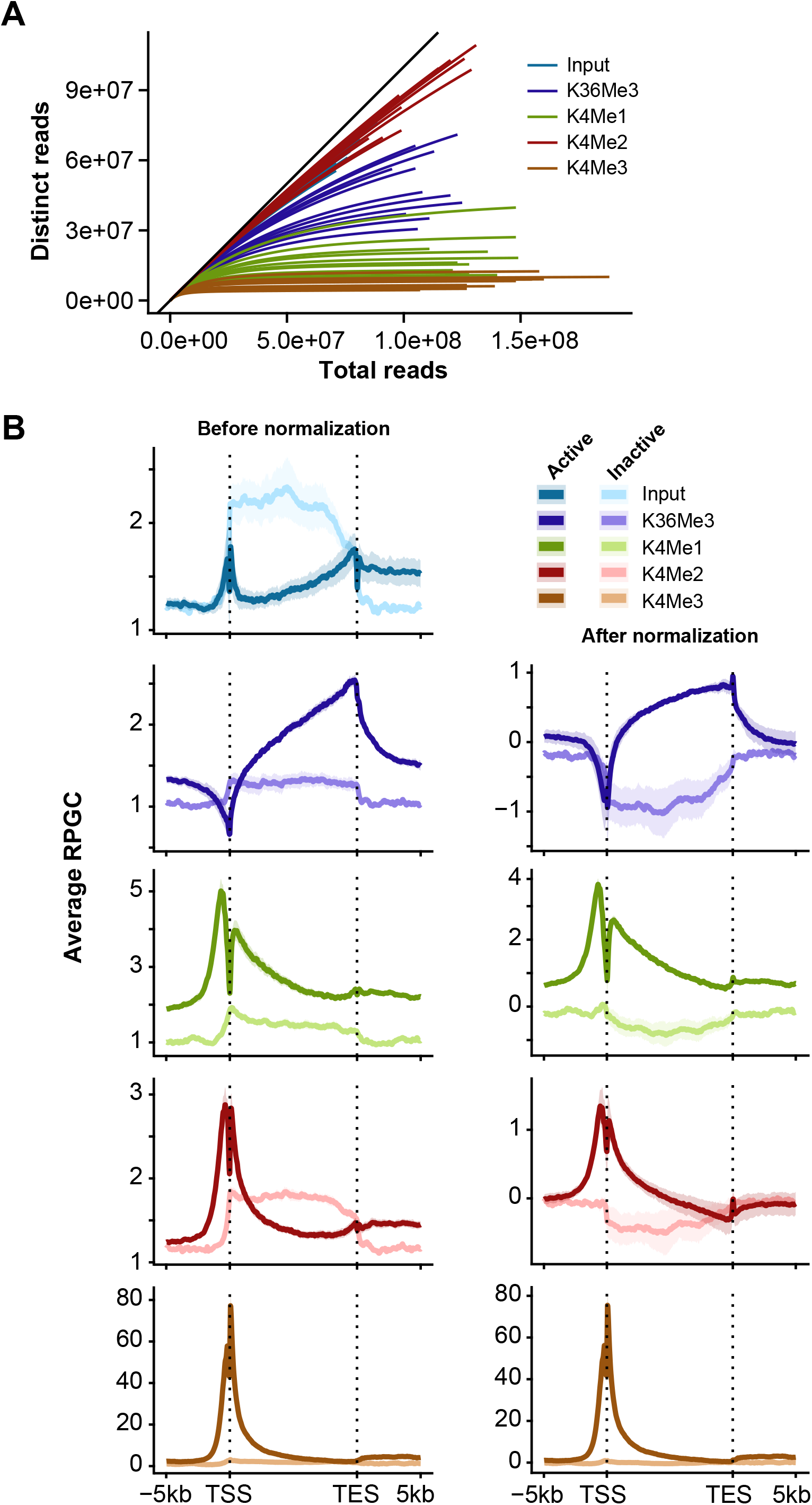
Optimization of cfChIP-seq. **(A)** Saturation curve of individual samples for input DNA and chromatin (H3K36Me3, H3K4Me1, H3K4Me2 and H3K4Me3) profile. **(B)** Distribution of cfChIP-seq signals on genes around the TSS and TES for with and without input normalization.

**Suppl Figure 2.**
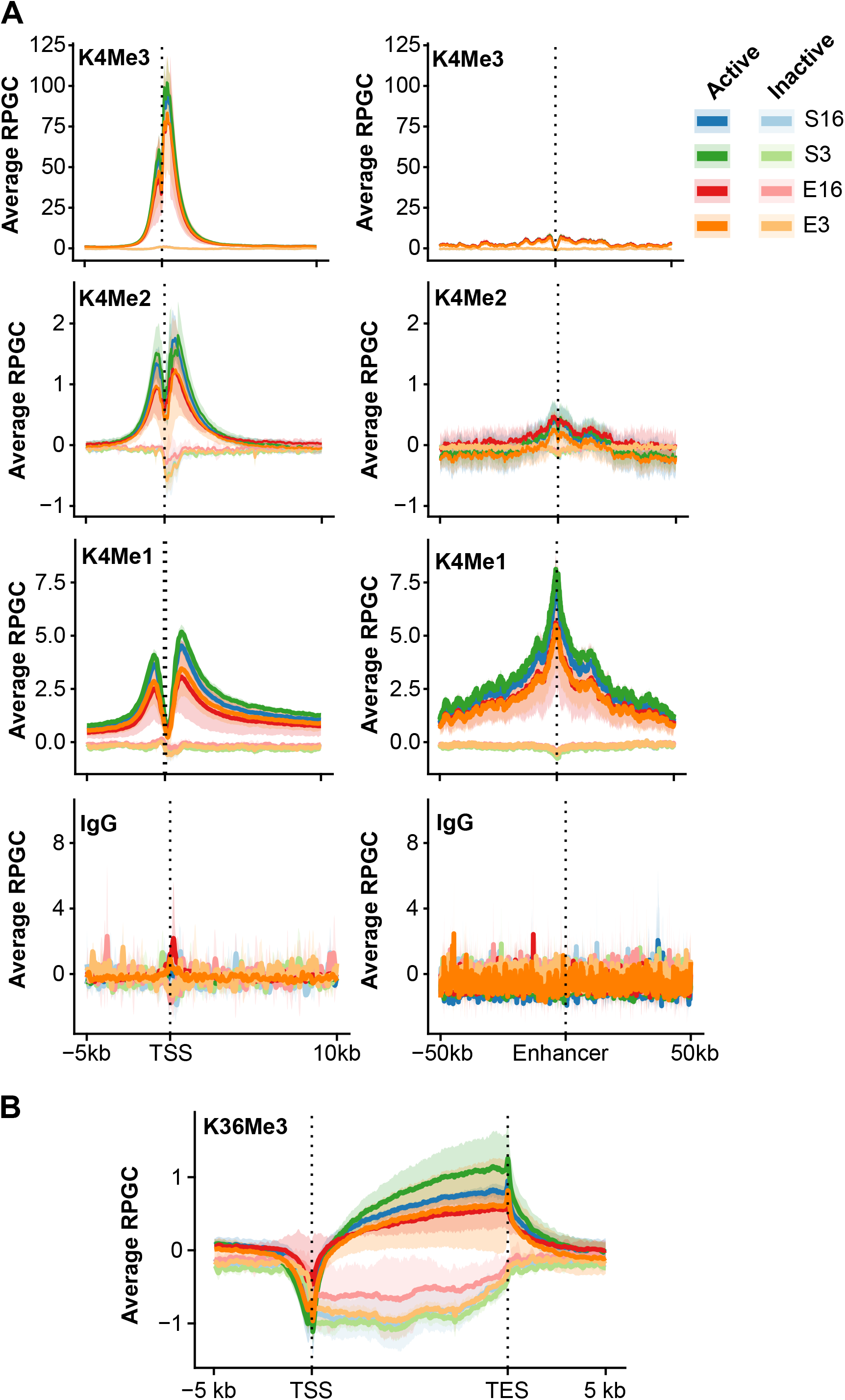
Comparable frequency distribution of chromatin profile among blood processing conditions. **(A)** Frequency distribution of H3K4me1, H3K4me2, H3K36me3, and IgG signals around the TSS in different processing conditions: E16, E3, S16, S3. **(B)** Frequency distribution of H3K36me3 across gene bodies in the different processing conditions.

**Suppl Figure 3.**
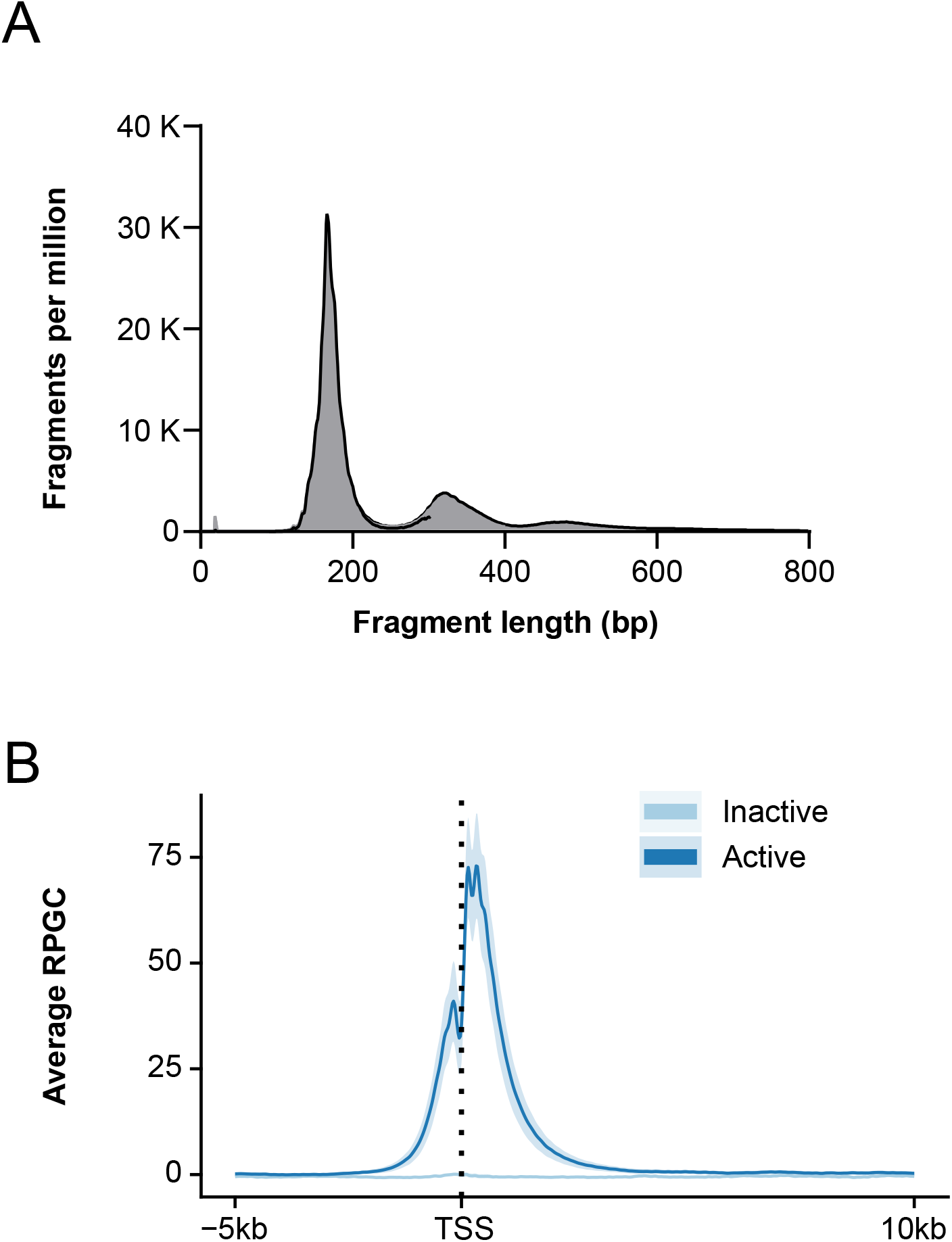
Profiles of the cell-free chromatin. **(A)** Length distribution of sequenced DNA fragments for heart transplant patients. **(B)** Frequency distribution for K4Me3 around the TSS site for heart transplant patients.

**Suppl Figure 4.**
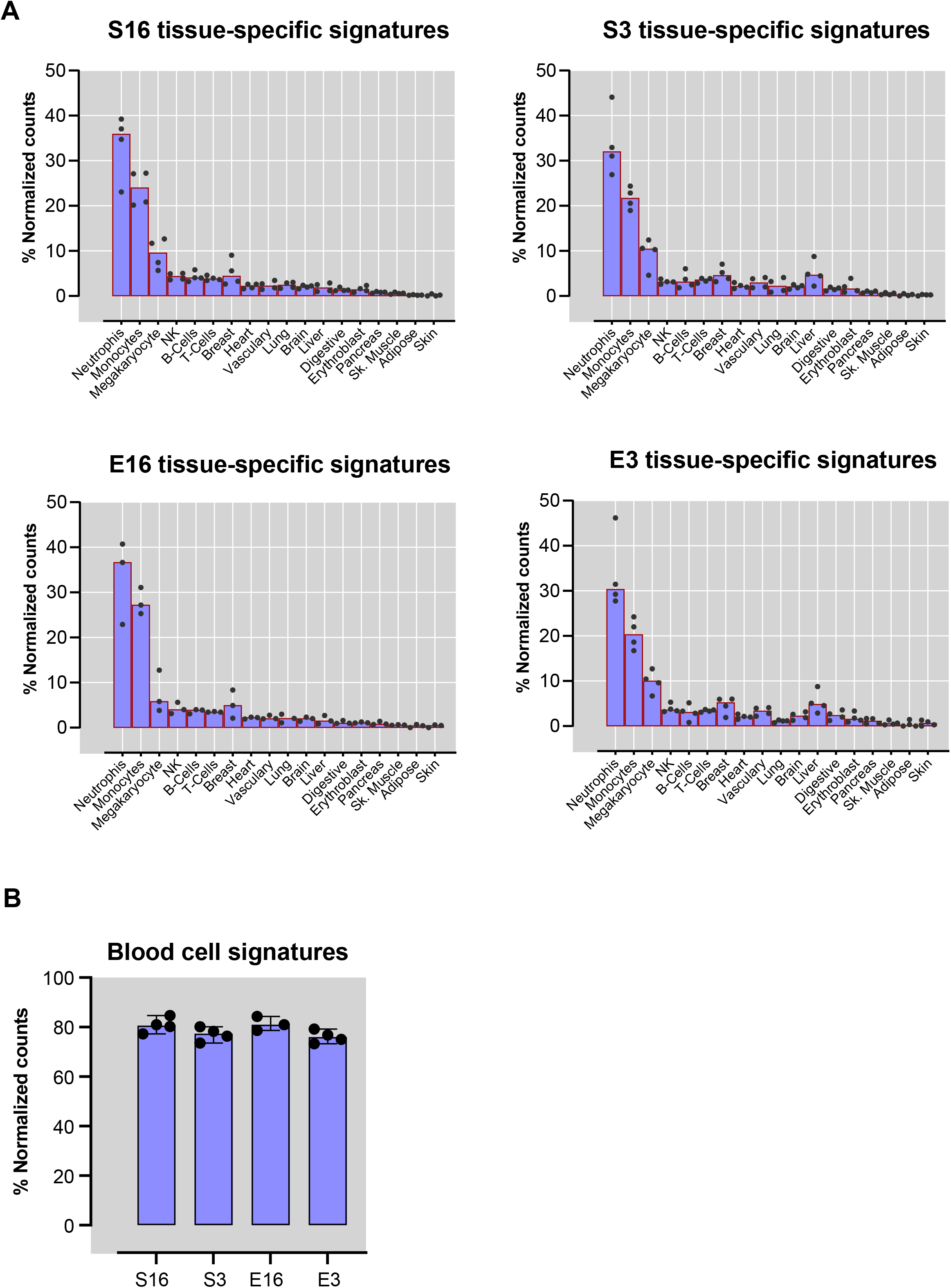
ChIP-seq tissue-specific signatures in different blood processing conditions. **(A)** Comparison of tissue-specific signatures among the four processing conditions (E16, E3, S16, S3). **(B)** The proportion of blood cells signatures between different conditions.

### Supplementary Table

Suppl. Table 1: Peaks, gene set with and without normalization for all four conditions

Suppl. Table 2: Housekeeping genes (also monocyte and neutrophil genes)

Suppl. Table 3: Number of differential genes by pathways

## Notes

### Competing Interest Statement

The authors have declared no competing interest.

## References

Agbor-Enoh S, Tunc I, De Vlaminck I, Fideli U, Davis A, Cuttin K, Bhatti K, Marishta A, Solomon MA, Jackson A. 2017. Applying rigor and reproducibility standards to assay donor-derived cell-free DNA as a non-invasive method for detection of acute rejection and graft injury after heart transplantation. The Journal of Heart and Lung Transplantation 36: 1004–1012.

Agbor-Enoh S, Wang Y, Tunc I, Jang MK, Davis A, De Vlaminck I, Luikart H, Shah PD, Timofte I, Brown AW et al. 2019. Donor-derived cell-free DNA predicts allograft failure and mortality after lung transplantation. EBioMedicine 40: 541–553.

Andargie TE, Tsuji N, Seifuddin F, Jang MK, Yuen PST, Kong H, Tunc I, Singh K, Charya A, Wilkins K. 2021. Cell-free DNA maps COVID-19 tissue injury and risk of death and can cause tissue injury. JCI insight 6.

Bonezi V, Genvigir FDV, Salgado PC, Felipe CR, Tedesco-Silva H, Jr., Medina-Pestana JO, Cerda A, Doi SQ, Hirata MH, Hirata RDC. 2020. Differential expression of genes related to calcineurin and mTOR signaling and regulatory miRNAs in peripheral blood from kidney recipients under tacrolimus-based therapy. Ann Transl Med 8: 1051.

Brusca SB, Elinoff JM, Zou Y, Jang MK, Kong H, Demirkale CY, Sun J, Seifuddin F, Pirooznia M, Valantine HA et al. 2022. Plasma Cell-Free DNA Predicts Survival and Maps Specific Sources of Injury in Pulmonary Arterial Hypertension. Circulation 146: 1033–1045.

Chandra S, Ehrlich KC, Lacey M, Baribault C, Ehrlich M. 2021. Epigenetics and expression of key genes associated with cardiac fibrosis: NLRP3, MMP2, MMP9, CCN2/CTGF and AGT. Epigenomics 13: 219–234.

De Weerdt A, Claeys KG, De Jonghe P, Ysebaert D, Chapelle T, Roeyen G, Jorens PG. 2008. Tacrolimus-related polyneuropathy: case report and review of the literature. Clin Neurol Neurosurg 110: 291–294.

Díaz B, González Vilchez F, Almenar L, Delgado JF, Manito N, Paniagua MJ, Crespo MG, Kaplinsky E, Pascual DA, Fernández-Yáñez J et al. 2007. Gastrointestinal complications in heart transplant patients: MITOS study. Transplant Proc 39: 2397–2400.

Duvvuri B, Lood C. 2019. Cell-Free DNA as a Biomarker in Autoimmune Rheumatic Diseases. Front Immunol 10: 502.

Fan S, Hu Y. 2022. Integrative analyses of biomarkers and pathways for heart failure. BMC Med Genomics 15: 72.

Fric J, Lim CX, Mertes A, Lee BT, Viganò E, Chen J, Zolezzi F, Poidinger M, Larbi A, Strobl H et al. 2014. Calcium and calcineurin-NFAT signaling regulate granulocyte-monocyte progenitor cell cycle via Flt3-L. Stem Cells 32: 3232–3244.

Jackson Chornenki NL, Coke R, Kwong AC, Dwivedi DJ, Xu MK, McDonald E, Marshall JC, Fox-Robichaud AE, Charbonney E, Liaw PC. 2019. Comparison of the source and prognostic utility of cfDNA in trauma and sepsis. Intensive Care Med Exp 7: 29.

Kechin A, Boyarskikh U, Kel A, Filipenko M. 2017. cutPrimers: A New Tool for Accurate Cutting of Primers from Reads of Targeted Next Generation Sequencing. J Comput Biol 24: 1138–1143.

Kondili M, Fust A, Preussner J, Kuenne C, Braun T, Looso M. 2017. UROPA: a tool for Universal RObust Peak Annotation. Sci Rep 7: 2593.

Kundaje A, Meuleman W, Ernst J, Bilenky M, Yen A, Heravi-Moussavi A, Kheradpour P, Zhang Z, Wang J, Ziller MJ et al. 2015. Integrative analysis of 111 reference human epigenomes. Nature 518: 317–330.

Li H, Durbin R. 2009. Fast and accurate short read alignment with Burrows-Wheeler transform. Bioinformatics 25: 1754–1760.

Li H, Handsaker B, Wysoker A, Fennell T, Ruan J, Homer N, Marth G, Abecasis G, Durbin R. 2009. The Sequence Alignment/Map format and SAMtools. Bioinformatics 25: 2078–2079.

Lindenfeld J, Miller GG, Shakar SF, Zolty R, Lowes BD, Wolfel EE, Mestroni L, Page RL, 2nd, Kobashigawa J. 2004. Drug therapy in the heart transplant recipient: part II: immunosuppressive drugs. Circulation 110: 3858–3865.

Lo YM, Zhang J, Leung TN, Lau TK, Chang AM, Hjelm NM. 1999. Rapid clearance of fetal DNA from maternal plasma. Am J Hum Genet 64: 218–224.

Marboe CC, Billingham M, Eisen H, Deng MC, Baron H, Mehra M, Hunt S, Wohlgemuth J, Mahmood I, Prentice J. 2005. Nodular endocardial infiltrates (Quilty lesions) cause significant variability in diagnosis of ISHLT Grade 2 and 3A rejection in cardiac allograft recipients. The Journal of heart and lung transplantation 24: S219–S226.

Moore JE, Purcaro MJ, Pratt HE, Epstein CB, Shoresh N, Adrian J, Kawli T, Davis CA, Dobin A, Kaul R et al. 2020. Expanded encyclopaedias of DNA elements in the human and mouse genomes. Nature 583: 699–710.

Moss J, Magenheim J, Neiman D, Zemmour H, Loyfer N, Korach A, Samet Y, Maoz M, Druid H, Arner P. 2018. Comprehensive human cell-type methylation atlas reveals origins of circulating cell-free DNA in health and disease. Nature communications 9: 1–12.

Nekrutenko A, Taylor J. 2012. Next-generation sequencing data interpretation: enhancing reproducibility and accessibility. Nat Rev Genet 13: 667–672.

Pruitt AA, Graus F, Rosenfeld MR. 2013. Neurological complications of solid organ transplantation. Neurohospitalist 3: 152–166.

Ramírez F, Ryan DP, Grüning B, Bhardwaj V, Kilpert F, Richter AS, Heyne S, Dündar F, Manke T. 2016. deepTools2: a next generation web server for deep-sequencing data analysis. Nucleic Acids Res 44: W160–165.

Richmond ME, Zangwill SD, Kindel SJ, Deshpande SR, Schroder JN, Bichell DP, Knecht KR, Mahle WT, Wigger MA, Gaglianello NA et al. 2020. Donor fraction cell-free DNA and rejection in adult and pediatric heart transplantation. J Heart Lung Transplant 39: 454–463.

Ross-Innes CS, Stark R, Teschendorff AE, Holmes KA, Ali HR, Dunning MJ, Brown GD, Gojis O, Ellis IO, Green AR. 2012. Differential oestrogen receptor binding is associated with clinical outcome in breast cancer. Nature 481: 389–393.

Sadeh R, Sharkia I, Fialkoff G, Rahat A, Gutin J, Chappleboim A, Nitzan M, Fox-Fisher I, Neiman D, Meler G et al. 2021. ChIP-seq of plasma cell-free nucleosomes identifies gene expression programs of the cells of origin. Nat Biotechnol 39: 586–598.

Şahintürk H, Yurtsever BM, Ersoy Ö, Kibaroğlu S, Zeyneloğlu P. 2021. Neurologic Complications in Heart Transplant Recipients Readmitted to the Intensive Care Unit. Cureus 13: e19425.

Tayyem O, Saraireh H, Al Hanayneh M, Stevenson HL. 2018. Heart transplant recipient with mycophenolate mofetil-induced colitis: the great imitator. BMJ Case Rep 2018.

Vorperian SK, Moufarrej MN, Quake SR. 2022. Cell types of origin of the cell-free transcriptome. Nat Biotechnol 40: 855–861.

Wang X, Bi Y, Xue L, Liao J, Chen X, Lu Y, Zhang Z, Wang J, Liu H, Yang H et al. 2015. The calcineurin-NFAT axis controls allograft immunity in myeloid-derived suppressor cells through reprogramming T cell differentiation. Mol Cell Biol 35: 598–609.

Zhang Y, Liu T, Meyer CA, Eeckhoute J, Johnson DS, Bernstein BE, Nusbaum C, Myers RM, Brown M, Li W et al. 2008. Model-based analysis of ChIP-Seq (MACS). Genome Biol 9: R137.

Zviran A, Schulman RC, Shah M, Hill STK, Deochand S, Khamnei CC, Maloney D, Patel K, Liao W, Widman AJ et al. 2020. Genome-wide cell-free DNA mutational integration enables ultra-sensitive cancer monitoring. Nat Med 26: 1114–1124.

